# Correcting values of DNA sequence similarity for errors in sequencing

**DOI:** 10.1101/237990

**Authors:** Timothy J. Hackmann

## Abstract

The similarity between two DNA sequences is one of the most important measures in bioinformatics, but errors introduced during sequencing make values of similarity lower than they should be. Here we develop a method to correct raw sequence similarity for sequencing errors and estimate the original sequence similarity. Our method is simple and consists of a single equation with terms for 1) raw sequence similarity and 2) error rates (e.g., from Phred quality scores). We show the importance of this correction for 16S ribosomal DNA sequences from bacterial communities, where 97% similarity is a frequent threshold for clustering sequences for analysis. At that threshold and typical error rate of 0.2%, correcting for error increases similarity by 0.36 percentage points. This result shows that, if uncorrected, sequencing error would increase similarity thresholds and generate false clusters for analysis. Our method could be used to adjust thresholds for cluster-based analyses. Alternatively, because it requires no clustering to correct sequence similarity, it could usher in a new age of analyzing ribosomal DNA sequences without clustering.

## Introduction

The similarity between two DNA sequences is one of the most important measures in bioinformatics. However, this measure is only as accurate as the sequences being compared. For example, two identical sequences will no longer be identical (have 100% similarity) after introducing sequencing errors. Errors pose a problem for determining the similarity of ribosomal DNA sequences from microbial communities, in particular, where 1) errors occur at high rates because of next generation sequencing and 2) biological variation is high and not easily distinguished from errors.

Many methods exist for handling sequencing errors in ribosomal DNA sequences, but most involve clustering. During clustering, similar sequences are grouped together, and one sequence (e.g., the most abundant) is presumed to be correct (representative of the cluster). This clustering is done by applying one similarity threshold (e.g., 97%)^1,2^, two thresholds sequentially (e.g., 99% and 97%)^3,4^, or more complex algorithms^5,6^. With 16S ribosomal DNA sequences, the nominal purpose of clustering at 97% similarity is to create discrete groups (operational taxonomic units) for analysis, but this clustering also masks sequence error.

Cluster-based methods are calibrated and evaluated with mock communities (mixtures of known species of microbes)^1–6^ (Fig. 1A). Clustering cannot be calibrated with real communities because species are not known. Because real communities differ from mock communities (e.g., by having more species), clustering with real communities may not be accurate (Fig. 1B). This represents a serious weakness of cluster-based methods.

**Fig. 1.**
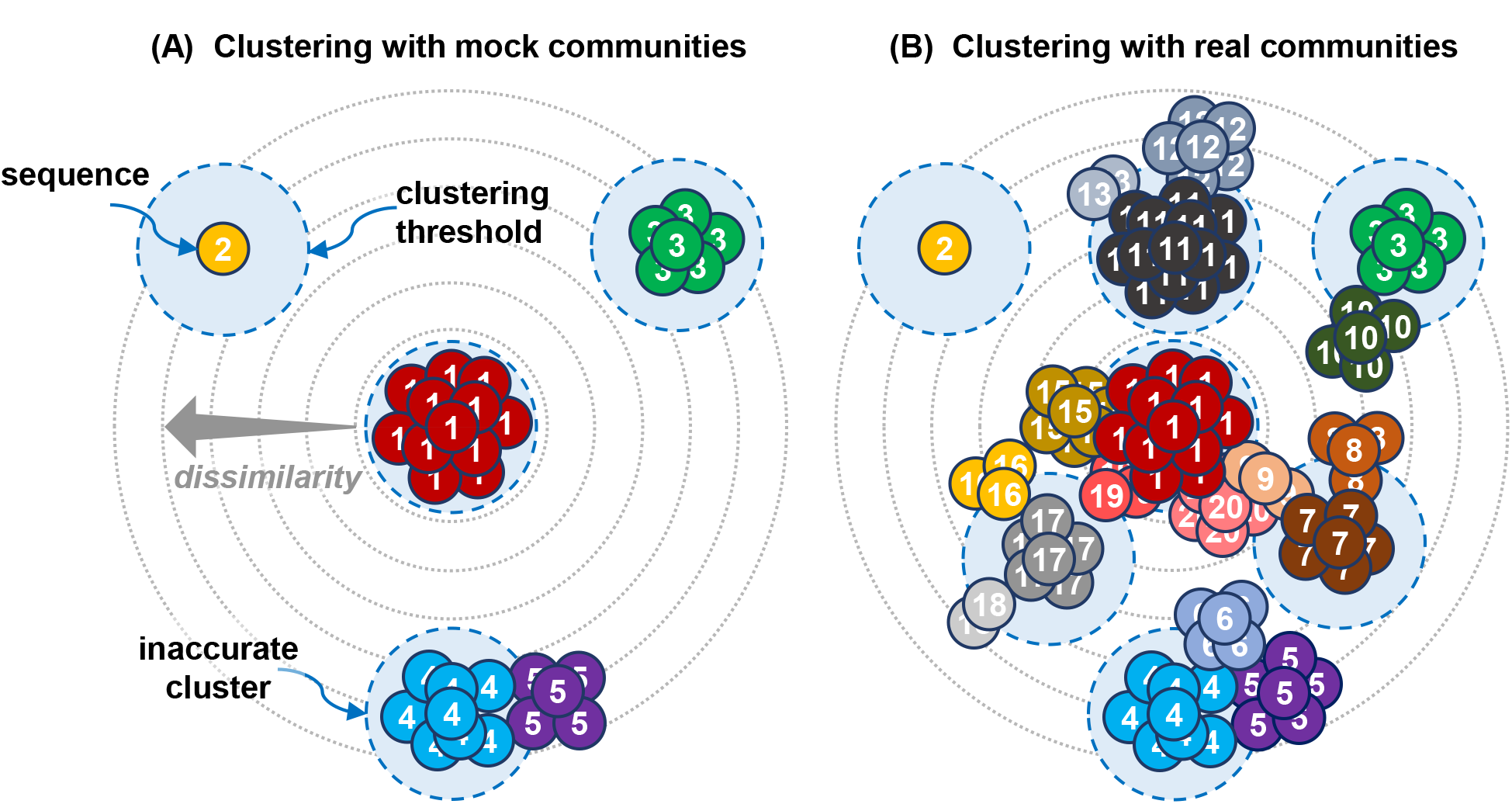
Cluster-based methods for handling sequence errors in ribosomal DNA sequences, illustrating their pitfalls. (A) Clustering thresholds are calibrated with mock communities of microbes to separate different sequences (species). Clusters are generally accurate (except for species 4 and 5 in this example). (B) Clustering cannot be calibrated with real communities because species are not known. Calibration from mock communities can be applied but may not be accurate (there are many clusters containing multiple species in this example). Sequences belonging to the same microbial species have the same color and number.

We present a simple method for handling errors in ribosomal and other DNA sequences. This method focuses on correcting sequence similarity (Fig. 2). This method permits analysis of ribosomal DNA sequence similarity and diversity without the pitfalls of clustering.

**Fig. 2.**
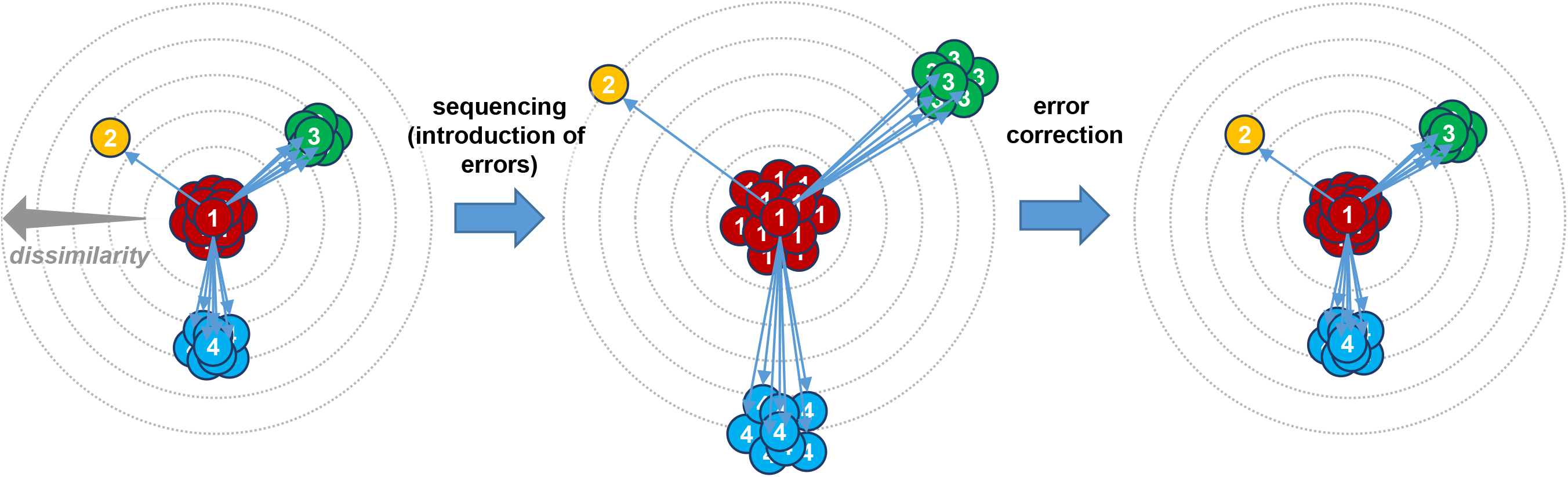
Our method for handling sequencing error in ribosomal DNA sequences; this method focuses on correcting sequence similarity. Sequences become more dissimilar after introducing sequencing errors, but similarities are corrected for error by application of eq. [13] or [14]. Dissimilarity is indicated by length of arrows. For simplicity, only a few arrows (those pertaining to the central sequence) are shown. Sequences belonging to the same microbial species have the same color and number.

## Results

We developed a method to correct the similarity of DNA sequences for sequencing errors. This method, which consists of a simple equation derived using Bayes’s theorem, provides an estimate of the original similarity (before introduction of errors). It requires only 1) raw sequence similarity (after introduction of sequencing errors) and 2) error rates (which can be calculated from Phred quality scores).

Using our method, we calculated the original sequence similarity, *P*(*S_b_*), expected at different values of raw similarity, *P*(*S_a_*), and sequencing error rates (Fig. 5). At 25% raw similarity (that expected for two unrelated sequences), original and raw similarity were equal. At >25% raw similarity, original similarity was higher than raw similarity, and the difference grew larger as raw similarity and the error rate increased.

Raw similarity of 97% is a threshold for clustering 16S ribosomal DNA sequences into the same operational taxonomic unit (see inset of Fig. 5). At 97% raw similarity [*P*(*S_a_*) = 0.97] and 1% error rate (*p_x_* = *p_y_* = 0.01), original similarity was 1.96 percentage points higher [*P*(*S_b_*) = 0.9896]. Even at a 0.2% error rate, the original similarity was still 0.39 percentage points higher [*P*(*S_b_*) = 0.9739]. An error rate of 1% corresponds to a typical value for the Illumina MiSeq platform and reads of the V4 region of 16S ribosomal DNA (after merging paired end reads with USEARCH)^5^. An error rate of 0.2% corresponds to those same reads subjected to stringent filtering (removal of reads with more than one expected error with USEARCH)^5^. In sum, original similarity was higher than raw similarity, even at low error rates, showing the importance in correcting similarity for sequence error.

To illustrate our method further, we applied it to simulated sequence reads (Fig. 6). We generated 15,000 pairs of reads with original similarity of 97% [(*S_b_*) = 0.97)], and then we introduced errors at a rate of 1%. Following introduction of errors, the similarity of the reads decreased (cf. Fig. 6A,B), as expected from Fig. 5. After applying our method for correcting similarity, the mean similarity increased back to its original value (cf. Fig. 6A,C). When we introduced errors at only 0.2%, results were similar, but the variance in Fig. 6E and F was less than in Fig. 6B and C. In sum, under the conditions of our simulation, our method provides an unbiased estimate of the original sequence similarity over a range of error rates. The benefit of reducing error was reducing variance in the estimate.

## Discussion

We developed a method to correct the similarity of two DNA sequences for errors in sequencing. The method is simple and requires only 1) raw (observed) sequence similarity and 2) error rates, which can be calculated from Phred quality scores. This method is important to any analysis using values of DNA sequence similarity. It is especially important for determining the similarity of ribosomal DNA sequences from microbial communities, owing to frequent errors that are hard to separate from biological variation.

Most methods for handling sequence errors with 16S ribosomal DNA sequences are based on clustering^1–6^. One sequence, usually the most abundant, is presumed correct (representative of the cluster). These methods can be evaluated with mock communities of bacteria, but there is no way to test their accuracy with real communities.

Our method requires no clustering and no presumption about which sequences are correct. Moreover, it shows that cluster-based methods are biased. With an error rate of 0.2%, clustering at an apparent (raw) similarity of 97% is in fact done at an actual (original) similarity of 97.39%. If error is not corrected, false clusters will be generated for analysis.

While our method could be used to adjust similarity thresholds in cluster-based analyses, it paves the way for cluster-free analysis. Analysis of diversity, for example, can be done using phylogenetic trees [see ref.^7, 8^], and tree construction requires only sequence similarities, not clusters. By correcting DNA similarities for sequence errors, our method removes a major bias in analysis of diversity. By accomplishing this without clustering, our method could usher in a new age of analyzing ribosomal DNA sequences directly, with no clustering step needed.

## Methods

### Overview

We will derive an equation to correct raw sequence similarity for sequencing errors. The derivation of this equation uses Bayes’s theorem and follows from work determining the probability of correct letters in paired-end sequence reads^5^.

Let *X* and *Y* be the letters of two sequences (Fig. 3). *X[_k_]* and *Y[_k_]* refer to a letter at a given position *k* within the sequences, and sequences have a total of *n* positions. Before sequencing (introduction of errors), the letters are *X_b_* and *Y*_b_. After sequencing (introduction of errors), some letters change, and the set of letters becomes *X_a_* and *Y_a_*. Let *S_b_* be letters in *X_b_* and *Y_b_* that are similar (when compared at a given position *k*), and *S_a_* are the letters similar between *X_a_* and *Y_a_*. The letters that are dissimilar are *D_a_* and *D_b_*. We partition *S_a_* as *S_a1_,* which originate from *S_b_*, and *S_a2_,* which originate from *D_b_*.

**Fig. 3.**
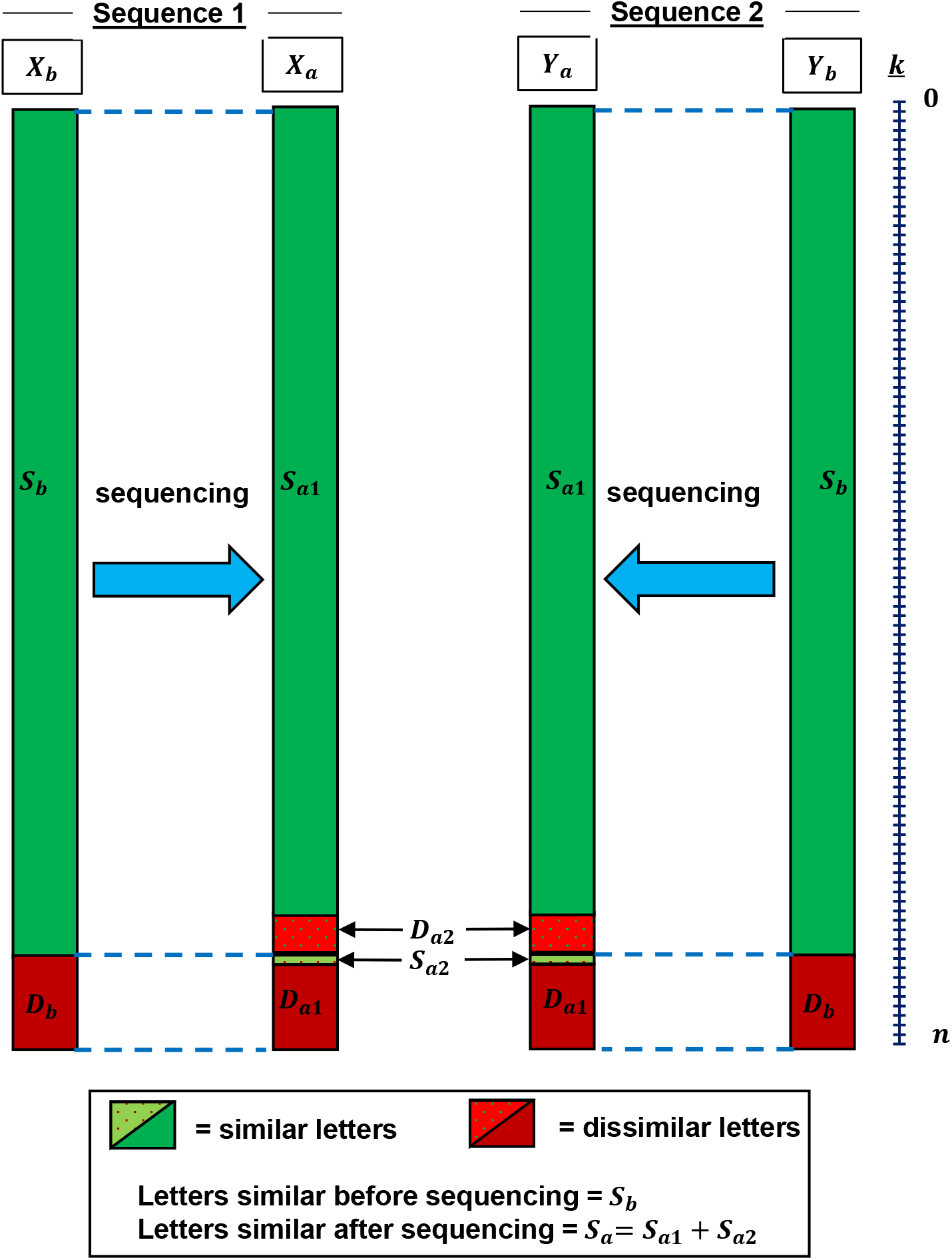
Two DNA sequences before and after sequencing, illustrating of terms in our equations for correcting sequence similarity. Letters in the two sequences are *X* and *Y*. Letters between *X* and *Y* that are similar are *S,* and letters dissimilar are *D.* The subscripts *a* and *b* (e.g., in *X_a_* and *X*_b_) refer to conditions before and after sequencing. Each letter has a position *k,* and there are a total of *n* positions. For illustration, letters in *S_b_* are grouped separately from letters in *D*_b_, though they would be interspersed in a real sequence. Letters in *S_a1_, S_a2_, D*_a1_, and *D_a2_* are grouped in the same way.

Our goal is to calculate the original similarity, *P*(*S_b_*), or similarity before introduction of errors. It is defined as *P*(*S_b_*) = *n_sb_/n*, where *n_sb_* is the number of positions in *S_b_*. We will calculate it from 1) the raw similarity, *P*(*S_a_*), or similarity after introduction of errors and 2) the error rates *p_x_* and *p_y_* (defined below).

### Similarity at a given position k

To estimate similarity of *X* and *Y* in total, we will first estimate the similarity at a given position in *X* and *Y. P*(*S_a_*[*_k_*]) is the raw similarity at position *k* and is either 0 (dissimilar) or 1 (similar). Following Fig. 4A, we can partition it as

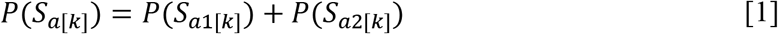

**Fig. 4.**
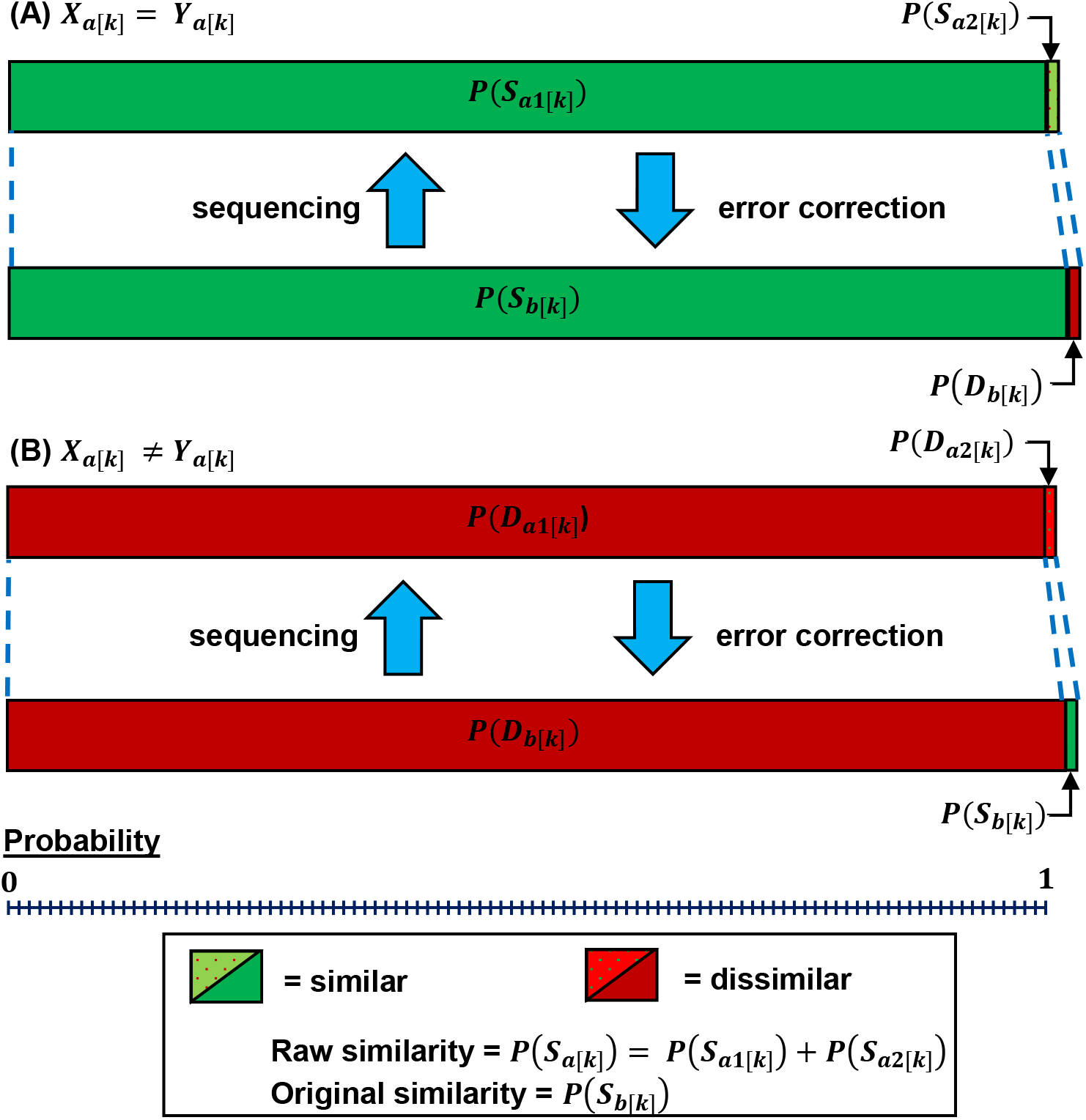
A given sequence position *(k)* before and after sequencing, illustrating of terms in our equations for correcting sequence similarity. (A) Condition where letters *X_a[k]_* and *Y_a[k]_* are similar [*P*(*S_a[k]_*) = 1]. (B) Condition where letters *X_a[k]_* and *Y_a[k]_* are dissimilar [*P*(*D_a[k]_*) = 1] In (A), *P*(*S_a2[k]_*) and *P*(*D_b[k]_*) are negative when *p_x[k]_* or *p_y[k]_* are positive, but they are depicted as positive for illustration. In (B), *P*(*D_a2[k]_*) and *P*(*S_b[k]_*) are negative when *p_x[k]_* or *p_y[k]_* is positive, but it is are depicted as positive for illustration.

By Bayes’s theorem

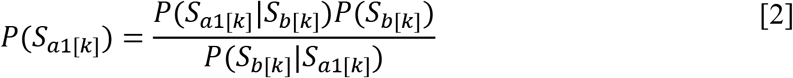

and

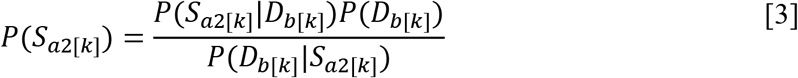

We note *P*(*S_b[k]_*) = 1 – *P*(*D_b[k]_*), substitute eq. [2] and [3] into [1], and solve for *P*(*S_b[k]_*) to give

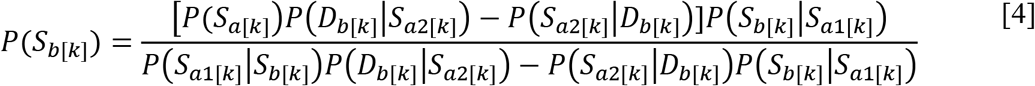

Next we find expressions for 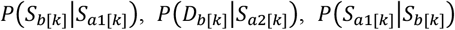, and 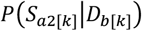. Because all *S*_*a*1[*k*]_ originate from *S_b[k]_* and all *S_a2[k]_* originate from *D_b[k]_* (Fig. 1),

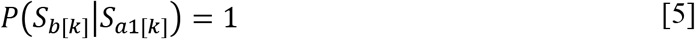

and

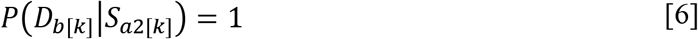

We partition 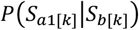 and 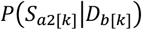

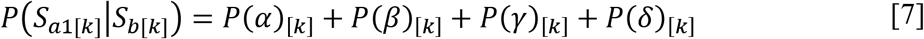

and

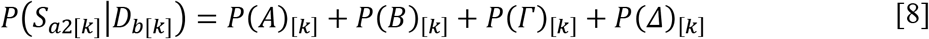

with terms defined in Table 1. For example, 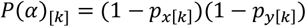 is the probability that neither *X_[k]_* nor *Y_[k]_* change (no errors were introduced) after sequencing, given the letters were similar before sequencing (i.e., *X_[k]_* and *Y_[k]_* belong to *S_b_*). The terms *p_x[k]_* and *p_y[k]_* are probabilities for change (sequencing error rate) for *X_[k]_* and *Y_[k]_*, respectively. We assume all changes (errors) are substitutions (not insertions or deletions), giving one of three equally probable outcomes per position per sequence. Phred quality scores *(Q)* can be used to calculate the error rates [e.g., 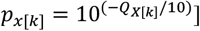.

**Table 1.**
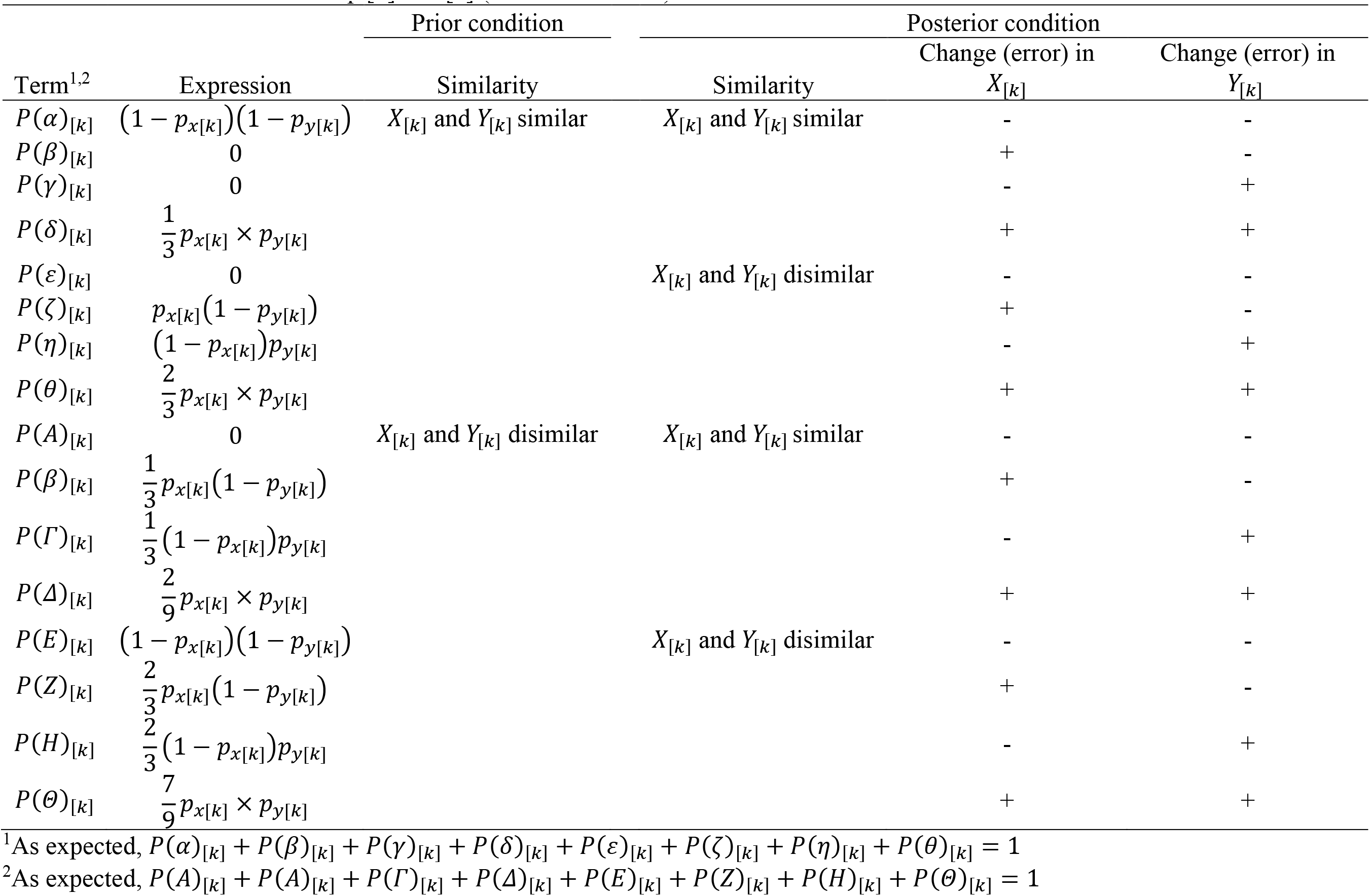
Definition of terms in eq. [7] and [8] (and related terms)

Substituting expressions for Table 1 into eq. [7] and [8] gives

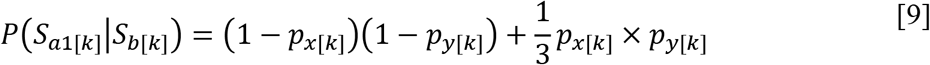

and

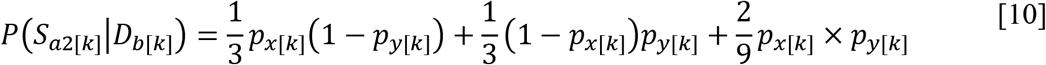

By substituting eq. [5], [6], [9], and [10] into eq. [4], we yield

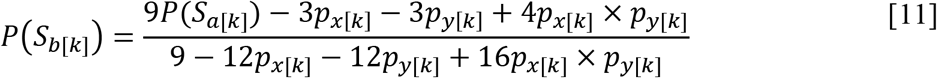

The derivation of eq. [11] is equivalent if we follow Fig. 4B and partition as *P*(*D_a[k]_*) as *p*(*D_a[k]_*) = *P*(*D_a1[k]_*) + *P*(*D_a2[k]_*) (not shown).

### Similarity across all n positions

To estimate similarity of *X* and *Y* in total, we average *P*(*S_b[k]_*) across all *n* positions

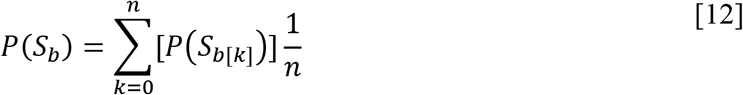

This approach assumes all changes (errors) occur independently (i.e., an error occurring at *k* = 0 does not change the probability of an error at *k* = 1). Eq. [12] can be expanding by substituting in eq. [11], giving

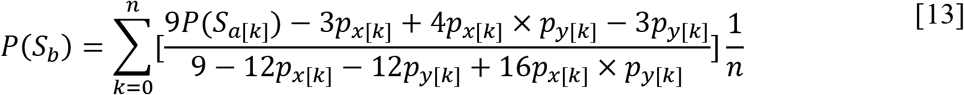

If we define *p_x_* as *p_x[k]_* averaged across *k* (and *p_y_* analogously), eq. [13] simplifies to

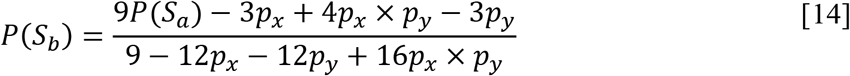

Eq. [13] or [14] permits us to calculate original sequence similarity, *P*(*S_b_*), given the raw sequence similarity, *P*(*S_a[k]_*), and error rates *p_x[k]_* and *p_y[k]_* [or *P*(*S_a_*), *p_x_* and *p_y_*]. These equations represent our method for correcting sequence similarity for sequencing error.

### Simulation of reads

We applied our method for correcting sequence similarity to sets of simulated reads. Reads with *n* = 300 positions each were generated. As above, errors were introduced under the modest assumptions that they 1) occur independently and 2) are in the form of substitutions. Other conditions of the simulation are specified in Fig. 6.

**Fig. 5.**
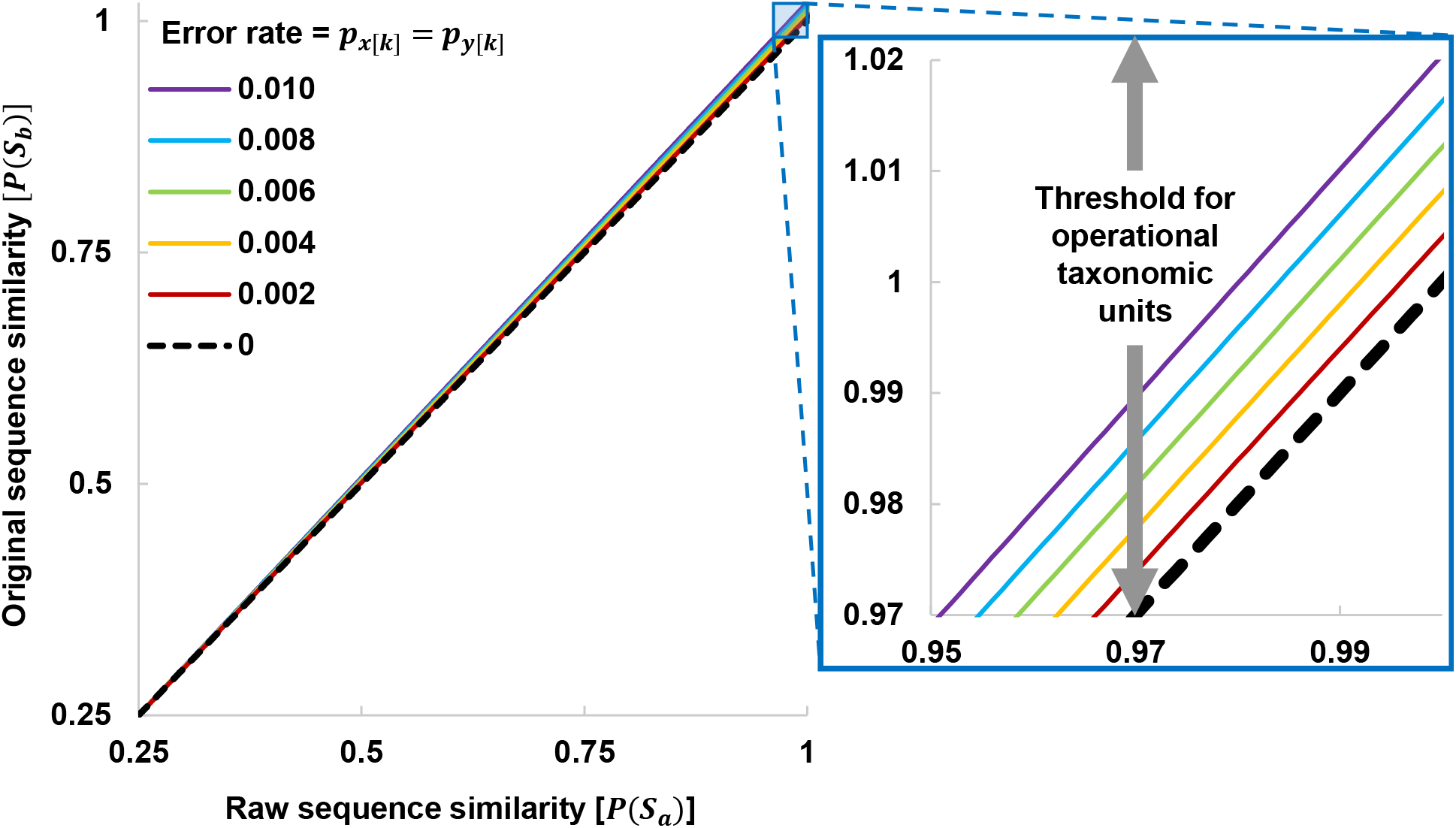
Comparison of raw sequence similarity [*P*(*S_a_*)] and original similarity [*P*(¾)]. Error rates vary from 0% (*p_x_* = *p_y_* = 0) to 1% (*p_x_* = *p_y_* = 0.01). Original similarity calculated from raw similarity using eq. [14].

**Fig. 6.**
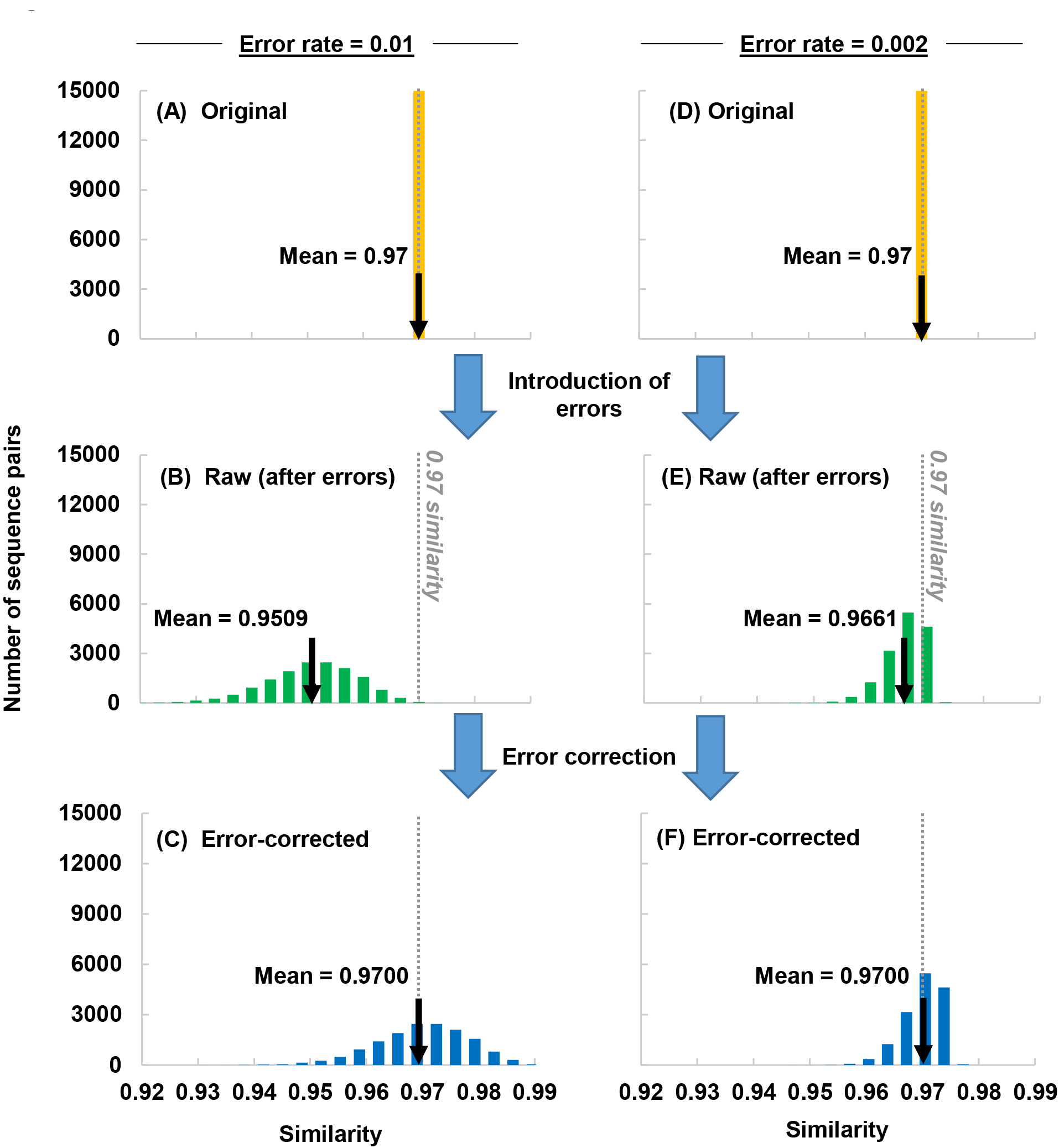
Our method for correcting sequencing similarity, as applied to 15,000 simulated sequence reads. (A) Reads before introduction of errors. Original similarity of reads was 97% [*P*(*S_b_*) = 0.97]. (B) Reads after introduction of errors at a rate of 1% (*p_x_* = *p_y_* = 0.01). (C) Reads after correcting similarity values in (B) with eq. [14]. Panels (D-F) are analogous, except error rate was 0.2% (*p_x_* = *p_y_* = 0.002).

## Acknowledgements

We thank J. Tao and S. Hackmann (University of Florida) for reviewing the manuscript. This work is supported by Agriculture and Food Research Initiative (AFRI) Competitive Grant no. 2017-67030-26589/project accession no. 1012177, Hatch Project accession no. 1002754, and Hatch Project accession no. 1002352 from the USDA National Institute of Food and Agriculture.

